# Implications of sample treatment on characterization of the riverine environmental metabolome

**DOI:** 10.1101/2021.09.02.458736

**Authors:** Amelia R. Nelson, Jason Toyoda, Rosalie K. Chu, Nikola Tolic, Vanessa A. Garayburu-Caruso, Casey M. Saup, Lupita Renteria, Jacqueline R. Wells, James C. Stegen, Michael J. Wilkins, Robert E. Danczak

## Abstract

High-resolution mass spectrometry techniques are widely used in the environmental sciences to characterize natural organic matter and, when utilizing these instruments, researchers must make multiple decisions regarding sample pre-treatment and the instrument ionization mode. To identify how these choices alter organic matter characterization and resulting conclusions, we analyzed a collection of 17 riverine samples from East River, CO (USA) under four PPL-based Solid Phase Extraction (SPE) treatment and electrospray ionization polarity (e.g., positive and negative) combinations: SPE (+), SPE (-), non-SPE (-), and non-SPE (+). The greatest number of formula assignments were achieved with SPE-treated samples due to the removal of compounds that could interfere with ionization. Furthermore, the SPE (-) treatment captured the most formulas across the widest chemical compound diversity. In addition to a reduced number of assigned formulas, the non-SPE datasets resulted in altered thermodynamic interpretations that could cascade into incomplete assumptions about the availability of organic matter pools for microbial heterotrophic respiration. Thus, we infer that the SPE (-) treatment is the best single method for characterizing environmental organic matter pools unless the focus is on lipid-like compounds, in which case we recommend a combination of SPE (-) and SPE (+) to adequately characterize these molecules.

**Synopsis:** We provide data-driven sample treatment and ionization mode recommendations to researchers who aim to use high-resolution mass spectrometry to characterize environmental organic matter.

**Abstract Art:** 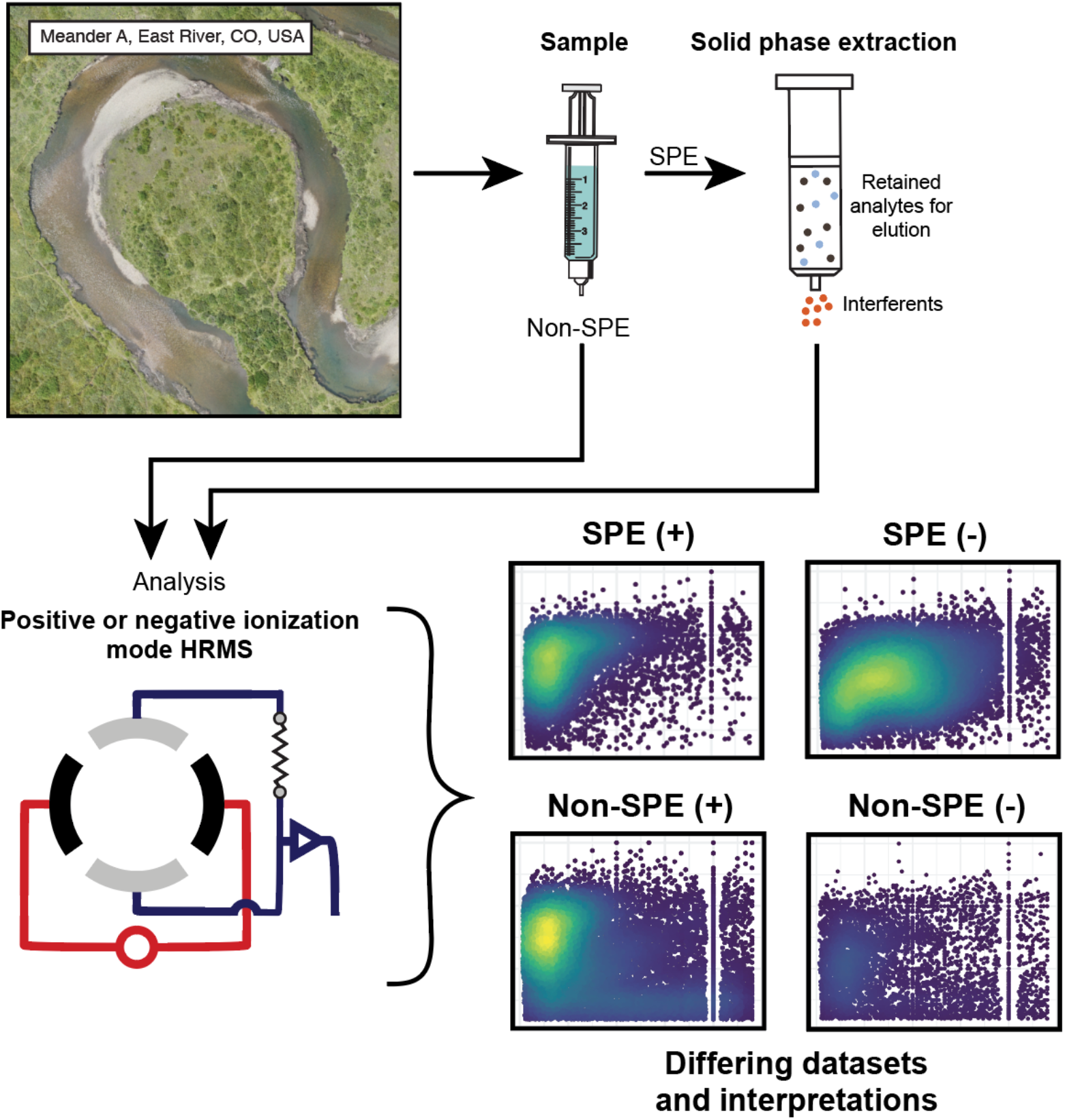

## Introduction

The development and increasing utilization of high-resolution mass spectrometry (HRMS) techniques has allowed scientists to characterize the molecular compounds that constitute natural organic matter^1^ (NOM). These approaches have been applied to samples across a diverse range of environments^2–5^, and have yielded new insights into NOM processing. More recently, attempts have been made to couple these high-resolution chemical analyses with microbiome data to directly link measurements of microbial function with changes in chemical species^6–8^. One of these approaches involves the analysis of dissolved organic matter (DOM) in a thermodynamic framework via which the nominal oxidation state of carbon (NOSC) can be calculated. Specifically, an empirical relationship between the NOSC and the standard molar Gibbs free energies of the oxidation half reactions of organic compounds^9^ allows researchers to quantify the thermodynamic favorability of DOM pools as electron donors for microbial heterotrophic respiration from HRMS data. One study using this approach demonstrated that aerobic respiration increased with increasing DOM thermodynamic favorability under carbon limiting conditions^6^. Similarly, a study of carbon dynamics within an anoxic aquifer revealed the preservation of DOM pools that yielded insufficient energy when coupled to the reduction of sulfate ^10^. Other recent studies taking a similar approach have further revealed relationships between DOM thermodynamics and various ecological measurements, including biogeochemical function and microbial community assembly^7,11–13^.

There are many pre-treatment and instrument options when utilizing HRMS techniques that could result in biasing the resulting dataset. Environmental samples can be analyzed without enrichment or treatment but are commonly pre-treated using PPL-based solid phase extraction (SPE) to concentrate DOM or remove species that may interfere with the ionization of organic compounds^14,15^ (e.g., salts). During SPE, DOM is retained on a sorbent then eluted using a solvent such as methanol. Polymer-based sorbents (e.g., PPL, C18) are commonly used during SPE due to their high extraction efficiency and representative character of retained DOM^16,17^. However, DOM recovery by SPE is incomplete and studies have shown that single-sorbent SPE methods can result in the preferential loss of specific compound classes such as organo-sulfur compounds^18^, aliphatic amines/amides, and tannin-like compounds with a high oxygen content^19,20^. Electrospray ionization (ESI) is a common ionization technique in HRMS because it can ionize a wide range of polar, hydrophilic molecules with diverse functional groups common in DOM, results in minimal-to-no fragmentation of DOM molecules (i.e., is a ‘soft’ ionization technique), and can be used in either positive or negative mode^21^. DOM can contain acidic functional groups that can be readily deprotonated in ESI (-) or basic functional groups (e.g., amines) that can be readily protonated and ionized efficiently in ESI (+). Thus, the chosen ESI mode can further govern the final HRMS DOM spectra and the chosen ESI mode has been found to result in different mass spectra for the same sample^22,23^. Ohno et al. (2016)^24^ found that ESI (+) was better at detecting aliphatic and carbohydrate-like molecules and that the ionization of these molecules was generally suppressed in samples rich in aromatics ionized in ESI (-) mode. The selected ESI mode can further impact downstream data through the generation of unintended adducts which can confound molecular formula assignment. For example, ESI (-) can result in the formation of chloride-containing ions due to salinity interference^16^. Finally, Hawkes et al. (2020)^25^ performed a cross-lab comparability study of HRMS data from different instruments to provide valuable metrics for future data benchmarking and further revealed patterns based upon ionization modes.

Although other studies have analyzed impacts of instrumentation, sample preparation, or ionization modes, no study has systematically examined the combined impacts of SPE and ionization mode choices or discussed changes to thermodynamic or microbial metabolic interpretations. Here, we analyze a set of 17 surface and pore water samples from the East River, CO, using four different Fourier transform ion cyclotron resonance mass spectrometry (FTICR-MS) methods (SPE or non-SPE, positive-ion or negative-ion ESI mode) with the goal of providing guidance to researchers on what method is most appropriate for their research question. We show that both sample pre-treatment and ESI mode choice influences the detection of different DOM molecules and can therefore bias the interpretation of microbial DOM processing within a given sample.

## Materials and Methods

### Sample collection

Surface and pore water samples (2 surface, 15 pore) were collected from the East River (CO, USA) on a 200-m reach encompassing one meander that lies within the U.S. Department of Energy-supported Lawrence Berkeley National Laboratory’s Watershed Science Focus Area (**Figure S1**). This combination of 17 samples were selected because they had representatives across all treatment types; see Nelson et al. (2019)^26^ and Saup et al. (2019)^27^ for more detailed information on sampling efforts and information on the remainder of the samples. Briefly, pore water samples were collected from their respective depths (**Table S1**) using a 0.6 cm diameter, stainless steel pore water sipper with a screen length of 5 cm attached to a syringe (MHE Products, MI, USA). One tubing volume of approximately 30 mL was discarded before sampling and sample was immediately filtered through 0.22 μm Sterivex filters (housings made of Eastar co-polyester; Massachusetts, USA). The 20 mL aliquot for the data presented here was filtered into a pre-combusted amber glass vial and immediately placed in a cooler on ice for transport back to the lab.

### Sample preparation and ESI-FTICR-MS data collection

Fourier Transform Ion Cyclotron Resonance Mass Spectrometry (FTICR-MS) was used to provide ultra-high resolution organic matter characterization. For the samples undergoing solid phase extraction, aqueous samples (NPOC 0.33-0.99 mg C/L) were acidified to pH 2 with 85% phosphoric acid and extracted with PPL cartridges (Bond Elut), following Dittmar et al. (2008)^16^. For those samples which were not extracted, 250 μL of sample were mixed with 500 μL of LC-MS grade MeOH. A 12 Tesla (12T) Bruker SolariX Fourier transform ion cyclotron resonance mass spectrometer (Bruker, Billerica, MA) located at the Environmental Molecular Sciences Laboratory in Richland, WA was used to collect high-resolution mass spectra of the organic matter found in each sample. Samples were directly injected into the instrument using a custom automated direct infusion cart that performed two offline blanks between each sample^28^. A Bruker SolariX electrospray ionization (ESI) source was used in positive and negative modes with applied voltages of +4.4kV and −4.2kV, respectively. Ion accumulation time was optimized between 50 and 80 ms. One hundred and forty-four transients were co-added into a 4MWord time domain (transient length of 1.1 s) with a spectral mass window of *m/z* 100-900, yielding a resolution at *m/z* 400. Spectra were internally recalibrated in the mass domain using homologous series separated by 14 Da (CH_2_ groups) (**File S1**). The mass measurement accuracy was typically within 1 ppm for singly charged ions across a broad m/z range (100 *m/z* - 900 *m/z*). Bruker Daltonics DataAnalysis (version 4.2) was used to convert mass spectra to a list of *m/z* values by applying the FTMS peak picking module with a signal-to-noise ratio (S/N) threshold set to 7 and absolute intensity threshold to the default value of 100. Formularity^29^ was used to assign chemical formulas based on exact mass, allowing a mass measurement error < 0.5 ppm and allowing for CHONS with the following restrictions: O>0 AND (N+S)<6 AND S<3 AND P=0. Formularity was also used to align peaks with a 0.5 ppm threshold.

The R package ftmsRanalysis^30,31^ was then used to remove peaks that either were outside the desired m/z range (150 m/z – 900 m/z) or had an isotopic signature, calculate nominal oxidation state of carbon (NOSC), assign putative compound classes^32^, and organize the data. We have included a table that describes the number of assigned peaks included after each filtering step (**Table S2**). Data published by Hawkes et al. (2020)^25^ was downloaded in R from https://github.com/BarrowResearchGroup/InterLabStudy. NOSC values were calculated and compound classes were assigned in order to compare broader datasets to the samples collected as part of this study.

### Thermodynamics Calculations

We calculated the average thermodynamic potential factor (F_T_) for the oxidation of average DOM pools coupled to the reduction of O_2_ and SO_4_^2-^ at standard conditions^9,33^. The F_T_ was calculated as follows:

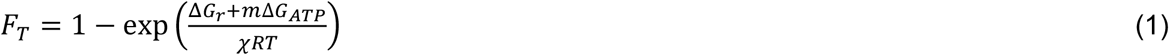

where ΔG_r_ is the Gibbs energy of the reaction, *m* is the number of moles of ATP synthesized per formula reaction (0.15 and 2 used for aerobic respiration and SO_4_^2-^ reduction, respectively^33^), ΔG_ATP_ denotes the Gibbs energy to synthesize ATP (50 kJ/mol used here^33^), χ is the average stoichiometric number for the reaction of interest, and R and T are the universal gas constant and temperature, respectively. Because our dataset represents average DOM pools, we chose to use an average value of 2 for the average stoichiometric number (χ).

### KEGG Mapping

In order to evaluate differences in potential biogeochemical interpretations, we assigned KEGG compound identifiers (CPD numbers) to observed molecular formulas by mapping them to the KEGG database using the provided REST API^34^. In order to avoid potential duplication of CPD numbers, only exact formula matches were considered. Using these CPD numbers, we identified corresponding pathways and qualitatively compared the most represented pathways across treatments. Web scraping was performed using the R package *rvest*^35^ and the KEGG mapping scripts are available on GitHub at https://github.com/danczakre/FTICR-Methods-Comparison.

### Statistics and Plot Generation

The statistics program R was used to perform all statistical analyses with the R package *ggplot2* used to generate all plots^36,37^. Comparisons across treatment groups were performed using Mann Whitney U tests (*wilcox.test)* in order to identify significant differences in compound classification, elemental composition, and average NOSC. An additional Kolmogorov-Smirnov test (*ks.test*) was used to evaluate difference in NOSC distributions within the Hawkes et al. dataset. Multivariate differences across treatment types were investigated using a nonmetric multidimensional scaling (NMDS) graph (*metaMDS*, vegan package v2.5-7) paired with a permuted analysis of variation (PERMANOVA; *adonis*, vegan package v2.5-7). All R scripts written to perform these analyses are available on GitHub at https://github.com/danczakre/FTICR-Methods-Comparison.

### Data Availability

All of the FTICR-MS data used throughout this study is available at the DOE’s data archive ESSDIVE at https://data.ess-dive.lbl.gov/view/doi:10.15485/1813303^38^.

## Results and discussion

### Molecular formula detection significantly varies based on sample preparation and ionization type

We evaluated 17 samples that were processed with or without SPE and analyzed in ESI (+) or ESI (-) mode to investigate the impact of method selection on FTICR-MS data. Results revealed clear differences in the collected data between methods (**Figure S2**). Considering the molecular formula count for each treatment, the SPE (-) treatment yielded the most assigned formulae (6218) while non-SPE (-) yielded the fewest (1257) (**Table S2**). More broadly, the SPE-treated samples contained greater molecular formula counts than non-SPE samples (**Figure 1**) and shared more common formulas (**Table S1)**. These results are expected because SPE is a common method to concentrate DOM in a sample and reduce the impact of salt during ionization^18,39^. By ensuring that carbon concentrations are higher, and that salts have a smaller impact on ionization efficiency, more molecular formula will be assigned. Differences in formula count between the two ionization modes are likely impacted by differences in the types of compounds ionized; specifically, river corridor organic matter is typically acidic and often has functional groups rich in oxygen which renders it more likely to be detected using negative mode^25,40^. The noted increase in molecular formula observed when comparing the non-SPE (-) vs. non-SPE (+), however, is likely the result of the SPE selecting for compounds more readily observed in negative mode. In other words, the molecular formulas observed under non-SPE conditions are more readily detectable in positive-mode or experience less ionization competition due to salinity. In these samples, conductivity (used as a proxy for salinity) ranged from 267.8 - 387.4 μS (**Table S1**). PPL-based solid-phase extraction can also lead to compositional shifts in the detected DOM given that some compound types have a higher affinity for the column (e.g., sulfur-containing, hydrophobic, low O/C compounds)^18,39^. The affinity of sulfur-containing compounds for the PPL-column is apparent in our dataset; SPE retention resulted in more assigned CHONS formulas in non-SPE in both negative and positive ionization mode (7.8% and 18.1%, respectively).

**Figure 1.**
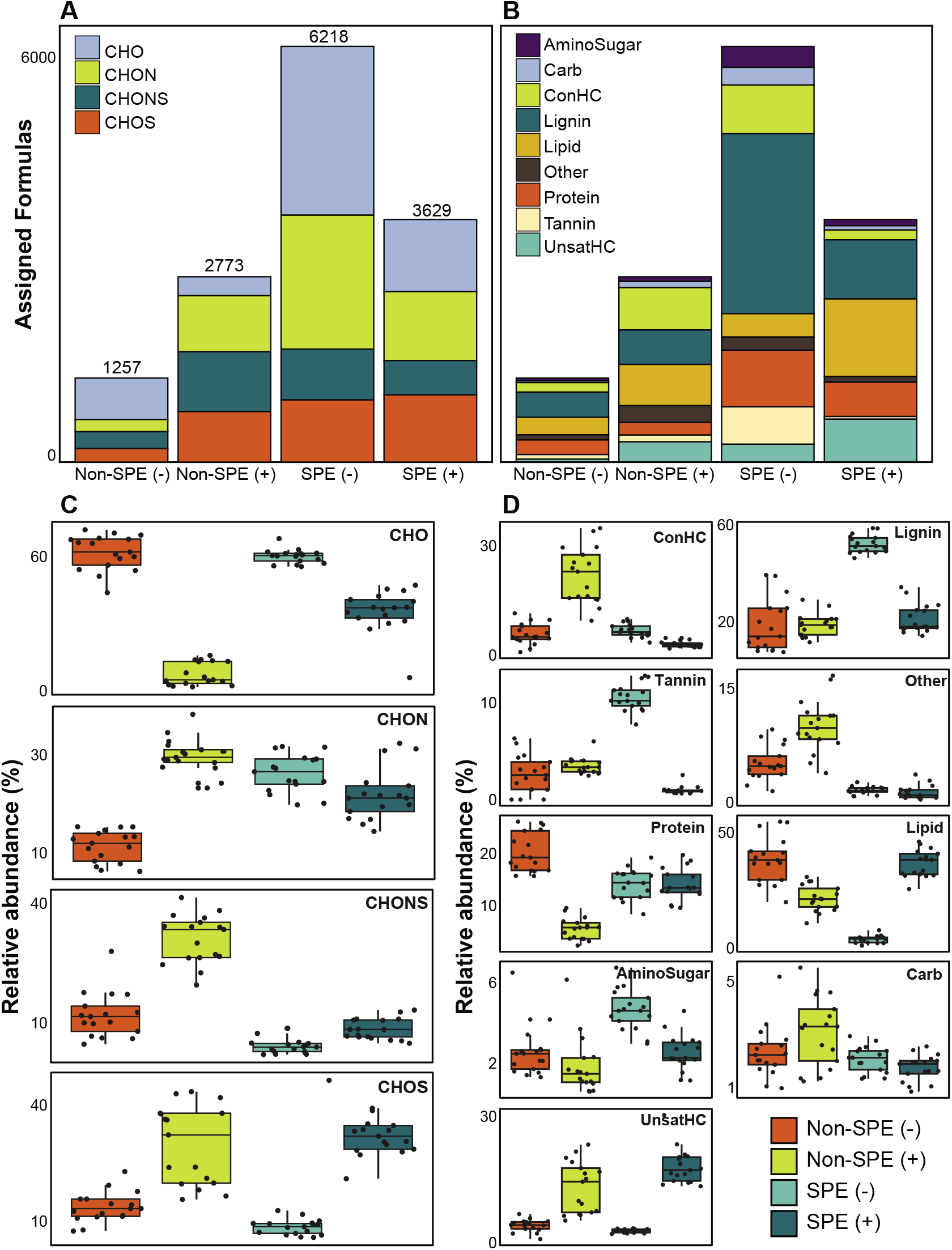
Classification of formulas assigned from each method by composition (**A**) and compound class (**B**). Numbers above bars (**A**) indicate the total number of detected and classified formulas for the sample set with each method. Trends are consistent in sample-resolved analyses of compositional (**C**) and compound class (**D**) variation between methods.

Broadly, we observed the largest variation the proportion of formulas assigned to be CHO-only, with values ranging from 10.2% in non-SPE (+) to 49.3% in non-SPE (-) and significant differences in proportion between each condition (Mann Whitney U p-value << 0.001) (**Figure 1a**). The SPE (-) samples had a similar proportion of CHO-only molecular formula as the non-SPE (-) samples (40.6% and 49.3%, respectively; Mann Whitney U p-value: 0.392), whereas the ESI (+) samples diverged in CHO-only formula proportions (10.2% and 29.7% in non-SPE and SPE, respectively). The large proportion of CHO-only formula in ESI (-) are consistent with observations that ESI (-) is primarily used to detect O-rich molecular formula^25,40^. CHON- and CHOS-containing formulas also demonstrated variability across treatment conditions overall (ranging from 14.4% to 32.2% and 14.9% to 27.7%, respectively; **Figure 1a**), while sample-resolved analyses reveal more complex patterns. ESI (+) samples, regardless of extraction method, consistently contained a higher proportion of CHOS-containing formula than ESI (-) samples (Mann Whitney U test p-value << 0.001). These patterns reflect the potential for ESI (+) to capture more compositionally complex spectra than ESI (-)^23^, whereas ESI (-) is more ideally suited for capturing CHO-only formula. Because environmental DOM is enriched by acidic, O-rich compounds^24,40,41^, the ESI (-) approach likely results in better characterization of the riverine DOM pool studied here.

Van Krevelen-based compound classes exhibited high variability across treatments as well (**Figure 1b**). While the non-SPE (+) treatment yielded more molecular formulas than non-SPE (-), both non-SPE ionization modes resulted in similar proportions of compound classes when all samples are considered together. The SPE (-) treatment generated the highest proportion of lignin- and tannin-like formulas (43.3% and 9%, respectively) while the SPE (+) treatment yielded a high proportion of lipid-like compounds (31.9%; **Figure 1b**). Sample-resolved analyses further showed these trends. Non-SPE (-) samples had a higher proportion of protein-like formulas than the other treatments (Mann Whitney U test p-value < 0.001) and non-SPE (+) had significantly greater representation of concentrated hydrocarbon-like formulas (Mann Whitney U test p-value << 0.001). Lignin-like and tannin-like formulas dominated SPE (-) samples (Mann Whitney p-value << 0.001) and the both non-SPE (-) and SPE (+) had increased lipid-like efficiency (Mann Whitney p-value < 0.001). Given that both the lignin- and tannin-like compound classes are characterized by higher O:C ratios (> 0.28 and > 0.65 respectively), we argue that these patterns reflect the enhanced capability the PPL cartridge to retain O-rich compounds^32,39^. This suggests that solid phase extraction has some combinatorial effect when used in conjunction with ESI (-) due to the elevated potential for ESI (-) to enrich for O-rich compounds, as we observed above.

### Thermodynamics vary substantially based upon extraction methodology and ionization mode

The NOSC for a given sample was used to evaluate the potential thermodynamic implications of the methodological differences. The NOSC metric can reveal the potential thermodynamic favorability of a carbon substrate (or bulk DOM pool favorability when averaged together), with higher NOSC values theoretically yielding a lower overall ΔG°_cox_ (i.e., more favorable) when coupled to the reduction of an electron acceptor^9^. We observed that each method yielded NOSC values significantly different from each other method (p < 0.001) and that compounds detected in the non-SPE (+) treatment had higher NOSC values overall while the average NOSC for detected compounds in the SPE (+) was significantly lower (**Figure 2**). Interestingly, the impacts of SPE on NOSC within each ionization mode are flipped; in ESI (+) mode, the SPE samples have significantly lower NOSC values whereas, in ESI (-) mode, the non-SPE samples have significantly lower NOSC values. We hypothesize that this could be due to potentially driven by the higher average proportion of lipid-like molecular formulas in both of these treatments, which have historically had lower NOSC values^10^. Thus, non-SPE (+) processing of these samples would have yielded DOM compounds considered less thermodynamically favorable in many biogeochemical analyses^6,10,42^.

**Figure 2.**
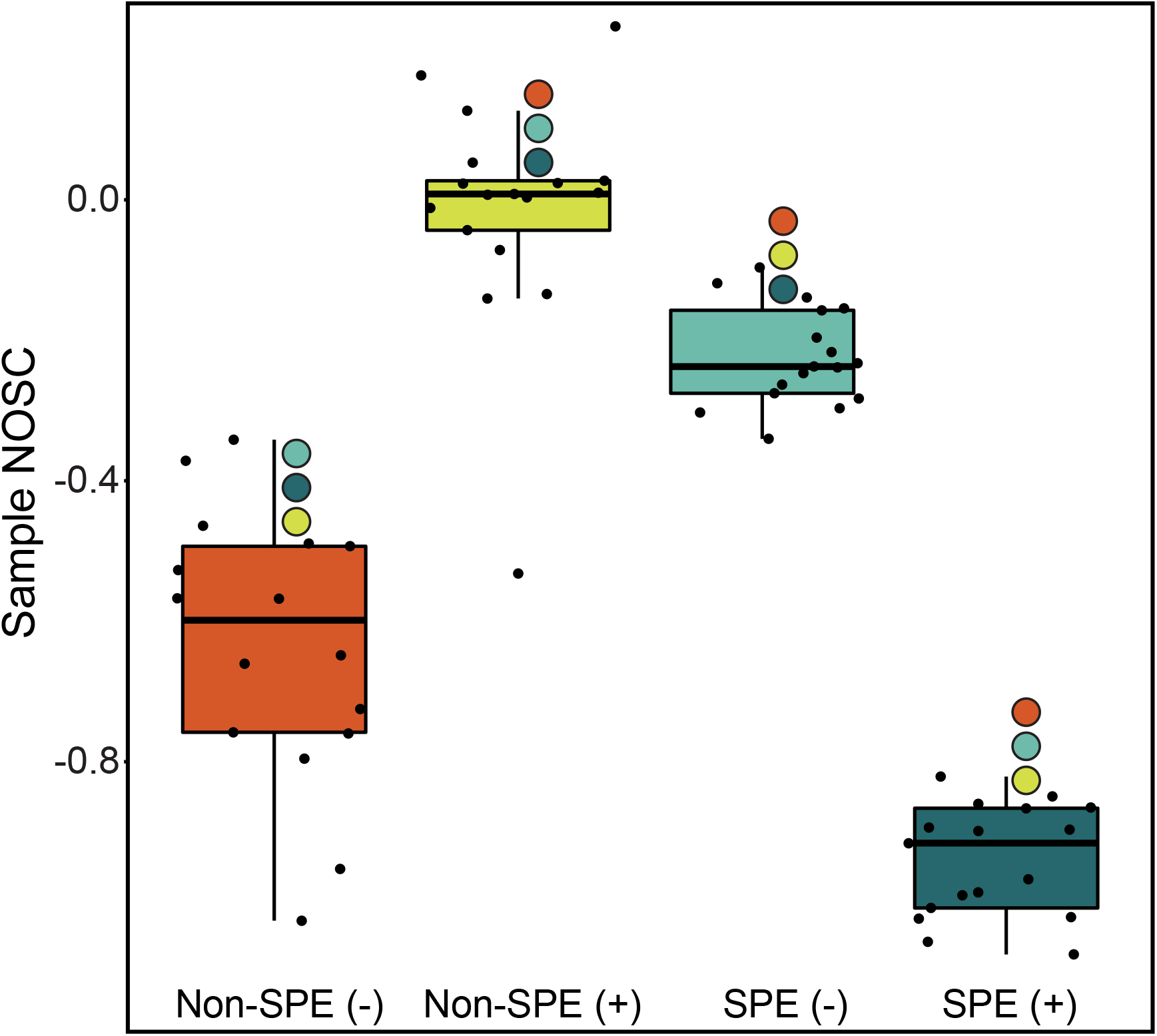
The distribution of sample mean NOSC values across all four FTICR-MS analysis methods. Colored circles indicate significant differences between methods.

To evaluate the transferability of observations derived from our dataset, we performed similar analyses on data previously utilized in Hawkes et al. (2020)^25^. In brief, these data consisted of four separate organic matter standards (ESFA: Elliot Soil Fulvic Acid, PLFA: Pony Lake Fulvic Acid, SRFA: Suwannee River Fulvic Acid, SRNOM: Suwannee River Natural Organic Matter) collected across international high-resolution mass spectrometry instruments (e.g., FTICR-MS, Orbitrap-MS) in both ESI (+) and ESI (-) modes. The fulvic acid samples were prepared by isolating DOM onto XAD-8 resin and the SRDOM sample was prepared using reverse osmosis. Across each standard alone and a composite of all standards, we observed consistently lower NOSC in data collected in positive mode than in negative mode (Wilcoxon Test & Kolmogorov-Smirnov Test p values << 0.001; **Figure S3**). The results help assert that the impact of ionization mode is similar across sample pre-treatment methods and sample types (e.g., ESFA is a soil standard vs. our freshwater samples).

To further explore the thermodynamic implications of the FTICR-MS methods, we calculated the average thermodynamic potential factor (F_T_) for the oxidation of the average DOM pools coupled to reduction of O_2_ and SO_4_^2-^ at standard conditions (**Figure 3**). Briefly, F_T_ is a dimensionless value derived from transition state theory that couples the rate of reaction to the Gibbs free energy of reaction. A value of 1 indicates that bioenergetic limitation is ignored, the reaction is more kinetically controlled, and, if there is substrate available, there will likely be a positive reaction rate until substrate is consumed^9,33^. Thus, F_T_ values are helpful for identifying differences in potential organic matter degradation rates between different environments or samples. Data resulting from the non-SPE (+) treatment consistently yielded the highest F_T_ values for reduction of both O_2_ and SO_4_^2-^, while the F_T_ values calculated from SPE (+) analyses were generally the lowest of the four treatment conditions (**Figure 3**). This indicates that the suite of compounds detected by non-SPE (+) would have the fewest thermodynamic constraints if utilized for sulfate reduction and that oxidation of the compounds detected by SPE (+) have the highest thermodynamic constraint when coupled to aerobic respiration. The compounds detected in SPE (-) consistently had high F_T_ values when coupled to aerobic oxidation and had a large range for sulfate reduction (0.02 - 0.51; **Figure 3**). As we observed with the NOSC values, we suggest that this is a result of non-SPE (-) and SPE (+) selecting for larger proportions of comparably thermodynamically unfavorable high H:C, low O:C formulas (e.g., lipid-like). As this approach is commonly coupled to FTICR-MS data^43^, it is important to understand how the pre-treatment and instrument method influences final F_T_ calculations and further biogeochemical interpretations.

**Figure 3.**
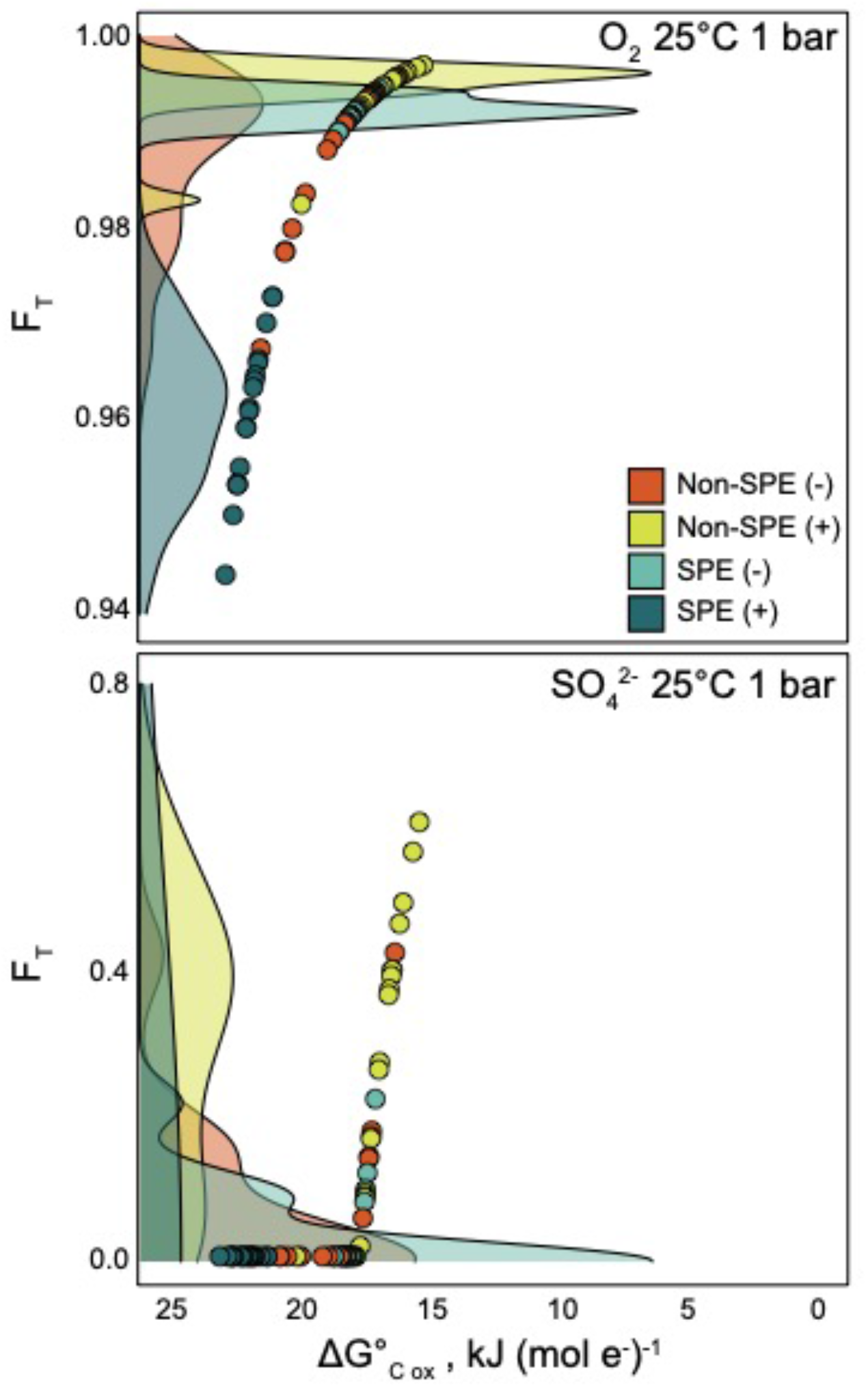
The Gibbs free energy of the oxidation of the average C pool plotted against the thermodynamic potential factor (F_T_) for the C oxidation coupled to O_2_ (top) or SO_4_^2-^ (bottom) reduction at standard conditions, colored by analysis method. Underlying colored density plots represent the distribution of F_T_ values across each method.

### Divergent ecosystem interpretations arise due to extraction methodology and ionization mode

We observed variability in elemental and lignin composition across ESI +/- and can leverage this to further evaluate the organic matter detected by each ionization mode. For example, lignin:N and C:N ratios have been used as an index of organic matter stability and as a means to estimate decomposition rate^44,45^. We calculated these widely used ratios for each of the datasets and found that the methods study here greatly impact these values (**Figure S4**). The non-SPE (+) dataset had a significantly lower C:N (here, organic C to organic N) than each other dataset and the SPE (-) dataset had a significantly higher lignin:N ratio due to the high detection of lignin-like compounds (**Figure 1**). This would lead to an interpretation that the DOM pool detected in non-SPE (+) has a higher residence time than those detected in the other methods. The higher lignin:N ratio in the SPE (-) dataset cascades into an interpretation where we would assume lower decomposition rates in this dataset relative to the others.

To identify whether the trends seen in the bulk dataset held for sample-resolved analyses, we chose a representative sample for direct comparison of selected measurements between the four methodologies (**Table 1**). A single representative sample was selected to focus specifically on the thermodynamic relationships across the treatment methodologies in one physical location, rather than broad scale biogeochemical processes explored elsewhere^26^. Overall, there were clear differences in the formula count, with SPE (-) still resulting in the most assigned formulas (1971). Additionally, despite variability in NOSC values across the sample treatments, measured DOM pools for this sample have few to no thermodynamic restraints for the reduction of O_2_ (i.e., F_T_ values close to 1 – **Figure 3a**). Conversely, these same treatment-derived differences in NOSC result in greater variability in the F_T_ parameter when oxidation of DOM is coupled to reduction of SO_4_^2-^. All the DOM pools detected using the SPE (+) and non-SPE (-) treatment yielded an F_T_ value of 0, indicating that sulfate reduction would be thermodynamically inhibited. In contrast, the non-SPE (+) produced the highest F_T_ when coupled to SO_4_^2-^ reduction, indicating fewer thermodynamic constraints. Lastly, similar to the bulk dataset analysis (**Figure S4**), the ratio of lignin-like compounds to N (lignin:N) was significantly higher in the SPE (-) dataset due to the large number of lignin-like compounds detected (**Figure 1B**) and the C:N ratio was still the lowest in the non-SPE (+). Thus, this example further illustrates how DOM treatment can influence our interpretation of likely redox reactions and potential decomposition rates occurring in a given sample.

**Table 1.**
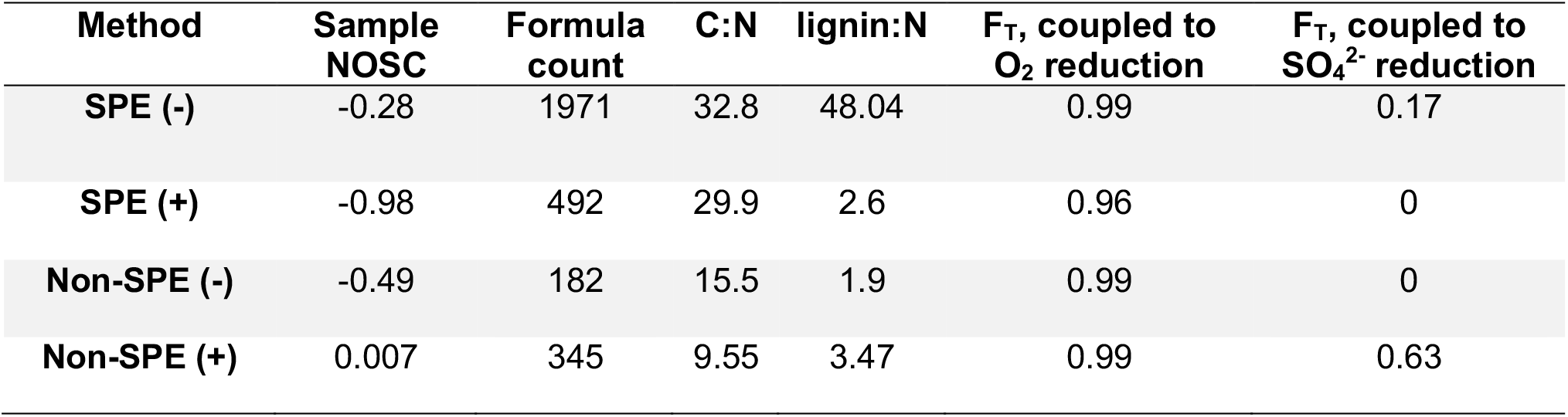
Results of one sample (23A, highlighted in Figure S1) analyzed using each of the four methods, indicating that the chosen method can result in varying conclusions and thermodynamic interpretations.

Given the observed patterns of unique and shared molecular formula across methodologies, extraction method and ionization mode have significant implications for biogeochemical modeling. In the case of recent multi-omics analysis, environmental metabolite data collected via high resolution MS can be incorporated into pathway models to understand carbon flux or organismal metabolism^46^. Differences in the types of formula detected during data collection could have profound impacts on inferred metabolic pathways. Given the low percentage of shared molecular formula across methods (**Table S3**), detected pathways may be divergent resulting in different predicted paths for carbon flux. Using the data presented within this manuscript, for example, we observe variation in the non-specific “Metabolic pathways” and “Biosynthesis of secondary metabolites” categories across the methods with SPE (-) having the highest absolute abundance and non-SPE (+) having the lowest (**Figure S7**). We also observe pathways unique to methods; for example, “Degradation of aromatic compounds” and “Flavonoid biosynthesis” were only present in SPE (-) data while we only saw “Limonene and pinene degradation” in SPE (+) (**Figure S7**). Other types of biogeochemical analyses, such as those relying on substrate-explicit modeling, will also be significantly impacted due to the shift in understood thermodynamic availability of each compound^12^. Specifically, the analysis proposed by Song et al. (2020)^12^ relies heavily on the predicted Gibb’s Free Energy of various carbon reactions to estimate reaction kinetics and stoichiometric coefficients of catabolic and anabolic reactions. Shifts in the average NOSC values will cascade into divergent predictions of biogeochemical rates.

## Conclusions

We achieved the highest number of assigned formulas (11,072) and representation from all molecular and van Krevelen-based compound classes when combining the four datasets (**Figure S5**). Although combining methods would most holistically characterize DOM pools, experimental design clearly depends on (1) instrument availability, (2) available sample volume, and (3) funding resources. We therefore provide some of the following recommendations to help researchers prioritize their desired outputs. Our analyses first confirm that SPE treatment yields the largest number of molecular formulas from environmental metabolomes and that ESI (-) captures the most formulas and is relatively sufficient for capturing broad chemical diversity (**Figure 1**). However, if lipid-like compounds (e.g., formulas with low O:C and high H:C) are the overall target, extracted samples run with ESI (+) would be a better recommendation. If the aim is a thermodynamic analysis of the DOM pools, SPE (+) appears to detect compounds with significantly lower NOSC values (**Figure 2**) than the other methods, potentially altering downstream thermodynamic predictions. We therefore recommend that SPE (-) or SPE (-) combined with SPE (+) is used for most applications. Non-SPE yields the lowest number of formulas and has the most limited applications due to salinity and other interferences. While it may be used if the laboratory has either time or funding constraints, we highlight the potential for saline interference in data capture. If samples are not treated with SPE, we recommend the use of ESI (+) instead of ESI (-) because it not only yields more formulas but also results in more formulas from each Van-Krevelen-based compound class, likely resulting in a more holistic characterization of the DOM pool (**Figure 1**). However, if the study is focusing on thermodynamics, ESI (-) detects compounds more representative of the DOM pool (**Figure 2**). These recommendations should be taken into consideration when performing experimental design.

## Abbreviations

SPE: solid phase extraction
HRMS: high-resolution mass spectrometry
NOM: natural organic matter
DOM: dissolved organic matter
NOSC: nominal oxidation state of carbon
ESI: electrospray ionization
FTICR-MS: Fourier transform ion cyclotron resonance mass spectrometry
NPOC: non-purgeable organic carbon

## Acknowledgments

A portion of this research was performed under the Facilities Integrating Collaborations for User Science (FICUS) program (award # 504279 to MJW) and used resources at the Environmental Molecular Sciences Laboratory (EMSL), which is a DOE Office of Science User Facility. The EMSL facility is sponsored by the Biological and Environmental Research (BER) program and operated under Contract No. DE-AC05-76RL01830. Another portion of this research was performed under DE-SC0016488 as part of the BER’s Subsurface Biogeochemical Research SFA funding of MJW. Sampling efforts were funded through a grant from the Geological Society of America to ARN. This research was also supported by the U.S. DOE-BER as part of BER’s River Corridor Research Program (RC). This contribution originates from the RC Scientific Focus Area (SFA) at the Pacific Northwest National Laboratory (PNNL). We would like to acknowledge the Worldwide Hydrobiogeochemical Observation Network for Dynamic River Systems (WHONDRS) for their support in data collection and analysis. We would also like to acknowledge William Kew for his assistance and guidance in the preparation in this manuscript.

## References

1. Hawkes, J. A., Dittmar, T., Patriarca, C., Tranvik, L. & Bergquist, J. Evaluation of the Orbitrap Mass Spectrometer for the Molecular Fingerprinting Analysis of Natural Dissolved Organic Matter. Anal. Chem. 88, 7698–7704 (2016).

2. Danczak, R. E. et al. Using metacommunity ecology to understand environmental metabolomes. Nat. Commun. 11, 6369 (2020).

3. Kellerman, A. M. et al. Fundamental drivers of dissolved organic matter composition across an Arctic effective precipitation gradient. Limnol. Oceanogr. 65, 1217–1234 (2020).

4. Kellerman, A. M., Dittmar, T., Kothawala, D. N. & Tranvik, L. J. Chemodiversity of dissolved organic matter in lakes driven by climate and hydrology. Nat. Commun. 5, 3804 (2014).

5. Tfaily, M. M., Hess, N. J., Koyama, A. & Evans, R. D. Elevated [CO2] changes soil organic matter composition and substrate diversity in an arid ecosystem. Geoderma 330, 1–8 (2018).

6. Garayburu-Caruso, V. A. et al. Carbon Limitation Leads to Thermodynamic Regulation of Aerobic Metabolism. Environ. Sci. Technol. Lett. 7, 517–524 (2020).

7. Graham, E. B. et al. Multi ‘omics comparison reveals metabolome biochemistry, not microbiome composition or gene expression, corresponds to elevated biogeochemical function in the hyporheic zone. Sci. Total Environ. 642, 742–753 (2018).

8. Graham, E. B. et al. Carbon Inputs From Riparian Vegetation Limit Oxidation of Physically Bound Organic Carbon Via Biochemical and Thermodynamic Processes. J. Geophys. Res. Biogeosciences 122, 3188–3205 (2017).

9. LaRowe, D. E. & Van Cappellen, P. Degradation of natural organic matter: A thermodynamic analysis. Geochim. Cosmochim. Acta 75, 2030–2042 (2011).

10. Boye, K. et al. Thermodynamically controlled preservation of organic carbon in floodplains. Nat. Geosci. 10, 415–419 (2017).

11. Sengupta, A. et al. Disturbance Triggers Non-Linear Microbe-Environment Feedbacks. Biogeosciences Discuss. 1–42 (2021). doi:10.1101/2020.09.30.314328

12. Song, H.-S. et al. Representing Organic Matter Thermodynamics in Biogeochemical Reactions via Substrate-Explicit Modeling. Front. Microbiol. 11, 1–16 (2020).

13. Stegen, J. C. et al. Influences of organic carbon speciation on hyporheic corridor biogeochemistry and microbial ecology James. Nat. Commun. 1–11 (2018). doi:10.1038/s41467-018-02922-9

14. Hertkorn, N., Harir, M., Koch, B. P., Michalke, B. & Schmitt-Kopplin, P. High-field NMR spectroscopy and FTICR mass spectrometry: Powerful discovery tools for the molecular level characterization of marine dissolved organic matter. Biogeosciences 10, 1583–1624 (2013).

15. King, R., Bonfiglio, R., Fernandez-Metzler, C., Miller-Stein, C. & Olah, T. Mechanistic investigation of ionization suppression in electrospray ionization. J. Am. Soc. Mass Spectrom. 11, 942–950 (2000).

16. Dittmar, T., Koch, B., Hertkorn, N. & Kattner, G. A simple and efficient method for the solid-phase extraction of dissolved organic matter (SPE-DOM) from seawater. Limnol. Oceanogr. Methods 6, 230–235 (2008).

17. Li, Y. et al. Proposed Guidelines for Solid Phase Extraction of Suwannee River Dissolved Organic Matter. Anal. Chem. 88, 6680–6688 (2016).

18. Tfaily, M. M., Hodgkins, S., Podgorski, D. C., Chanton, J. P. & Cooper, W. T. Comparison of dialysis and solid-phase extraction for isolation and concentration of dissolved organic matter prior to Fourier transform ion cyclotron resonance mass spectrometry. Anal. Bioanal. Chem. 404, 447–457 (2012).

19. Perminova, I. V. et al. Molecular Mapping of Sorbent Selectivities with Respect to Isolation of Arctic Dissolved Organic Matter as Measured by Fourier Transform Mass Spectrometry. Environ. Sci. Technol. 48, 7461–7468 (2014).

20. Sleighter, R. L. & Hatcher, P. G. Molecular characterization of dissolved organic matter (DOM) along a river to ocean transect of the lower Chesapeake Bay by ultrahigh resolution electrospray ionization Fourier transform ion cyclotron resonance mass spectrometry. Mar. Chem. 110, 140–152 (2008).

21. Sleighter, R. L. & Hatcher, P. G. The application of electrospray ionization coupled to ultrahigh resolution mass spectrometry for the molecular characterization of natural organic matter. J. Mass Spectrom. 42, 559–574 (2007).

22. Brown, T. L. & Rice, J. A. Effect of experimental parameters on the ESI FT-ICR mass spectrum of fulvic acid. Anal. Chem. 72, 384–390 (2000).

23. Rostad, C. E. & Leenheer, J. A. Factors that affect molecular weight distribution of Suwannee river fulvic acid as determined by electrospray ionization/mass spectrometry. Anal. Chim. Acta 523, 269–278 (2004).

24. Ohno, T., Sleighter, R. L. & Hatcher, P. G. Comparative study of organic matter chemical characterization using negative and positive mode electrospray ionization ultrahigh-resolution mass spectrometry. Anal. Bioanal. Chem. 408, 2497–2504 (2016).

25. Hawkes, J. A. et al. An international laboratory comparison of dissolved organic matter composition by high resolution mass spectrometry: Are we getting the same answer? Limnol. Oceanogr. Methods 18, 235–258 (2020).

26. Nelson, A. R. et al. Heterogeneity in Hyporheic Flow, Pore Water Chemistry, and Microbial Community Composition in an Alpine Streambed. J. Geophys. Res. Biogeosciences 2019JG005226 (2019). doi:10.1029/2019JG005226

27. Saup, C. M. et al. Hyporheic Zone Microbiome Assembly Is Linked to Dynamic Water Mixing Patterns in Snowmelt-Dominated Headwater Catchments. J. Geophys. Res. Biogeosciences 124, 3269–3280 (2019).

28. Orton, D. J. et al. A Customizable Flow Injection System for Automated, High Throughput, and Time Sensitive Ion Mobility Spectrometry and Mass Spectrometry Measurements. Anal. Chem. 90, 737–744 (2018).

29. Tolić, N. et al. Formularity: Software for Automated Formula Assignment of Natural and Other Organic Matter from Ultrahigh-Resolution Mass Spectra. Anal. Chem. 89, 12659–12665 (2017).

30. Bramer, L. M. & White, A. ftmsRanalysis: Analysis and visualization tools for FT-MS data. R package version 1.0.0. (2019).

31. Bramer, L. M. et al. ftmsRanalysis: An R package for exploratory data analysis and interactive visualization of FT-MS data. PLOS Comput. Biol. 16, e1007654 (2020).

32. Kim, S., Kramer, R. W. & Hatcher, P. G. Graphical Method for Analysis of Ultrahigh-Resolution Broadband Mass Spectra of Natural Organic Matter, the Van Krevelen Diagram. Anal. Chem. 75, 5336–5344 (2003).

33. Jin, Q. & Bethke, C. M. Predicting the rate of microbial respiration in geochemical environments. Geochim. Cosmochim. Acta 69, 1133–1143 (2005).

34. Kanehisa, M. & Goto, S. KEGG: Kyoto Encyclopedia of Genes and Genomes. Nucleic Acids Res. 28, 27–30 (2000).

35. Wickham, H. rvest: Easily Harvest (Scrape) Web Pages. (2020).

36. Team, R. C. R: A language and environment for statistical computing. R Found. Stat. Comput. Vienna, Austria (2021).

37. Wickham, H. ggplot2: Elegant Graphics for Data Analysis. (2016).

38. Nelson, A. R. et al. East River Surface and Pore Water FTICR-MS Data Associated with ‘Implications of sample treatment on characterization of the riverine environmental metabolome’. Seas. Control. Dyn. hyporheic Zo. redox Biogeochem. (2021). doi:10.15485/1813303

39. Raeke, J., Lechtenfeld, O. J., Wagner, M., Herzsprung, P. & Reemtsma, T. Selectivity of solid phase extraction of freshwater dissolved organic matter and its effect on ultrahigh resolution mass spectra. Environ. Sci. Process. Impacts 18, 918–927 (2016).

40. Perdue, E. M. & Ritchie, J. D. Dissolved Organic Matter in Freshwaters. in Treatise on Geochemistry 5–9, 273–318 (Elsevier, 2003).

41. Rostad, C. E. & Leenheer, J. A. Factors that affect molecular weight distribution of Suwannee river fulvic acid as determined by electrospray ionization/mass spectrometry. Anal. Chim. Acta 523, 269–278 (2004).

42. Danczak, R. E. et al. Ecological theory applied to environmental metabolomes reveals compositional divergence despite conserved molecular properties. Sci. Total Environ. 788, 147409 (2021).

43. Keiluweit, M., Wanzek, T., Kleber, M., Nico, P. & Fendorf, S. Anaerobic microsites have an unaccounted role in soil carbon stabilization. Nat. Commun. 8, 1771 (2017).

44. Melillo, J. M., Aber, J. D. & Muratore, J. F. Nitrogen and lignin control of hardwood leaf litter decomposition dynamics. Ecology 63, 621–626 (1982).

45. Parnas, H. Model for decomposition of organic material by microorganisms. Soil Biol. Biochem. 7, 161–169 (1975).

46. Kessell, A. K., McCullough, H. C., Auchtung, J. M., Bernstein, H. C. & Song, H.-S. Predictive interactome modeling for precision microbiome engineering. Curr. Opin. Chem. Eng. 30, 77–85 (2020).

