## Supplemental Information for "Implications of sample treatment on characterization of the riverine environmental metabolome"

### Table of contents

| | |
|---|---|
| Supplementary Figures 1-7 | 2-7 |
| Supplementary Table 1 | 8 |
| Supplementary Tables 2,3 | 9 |

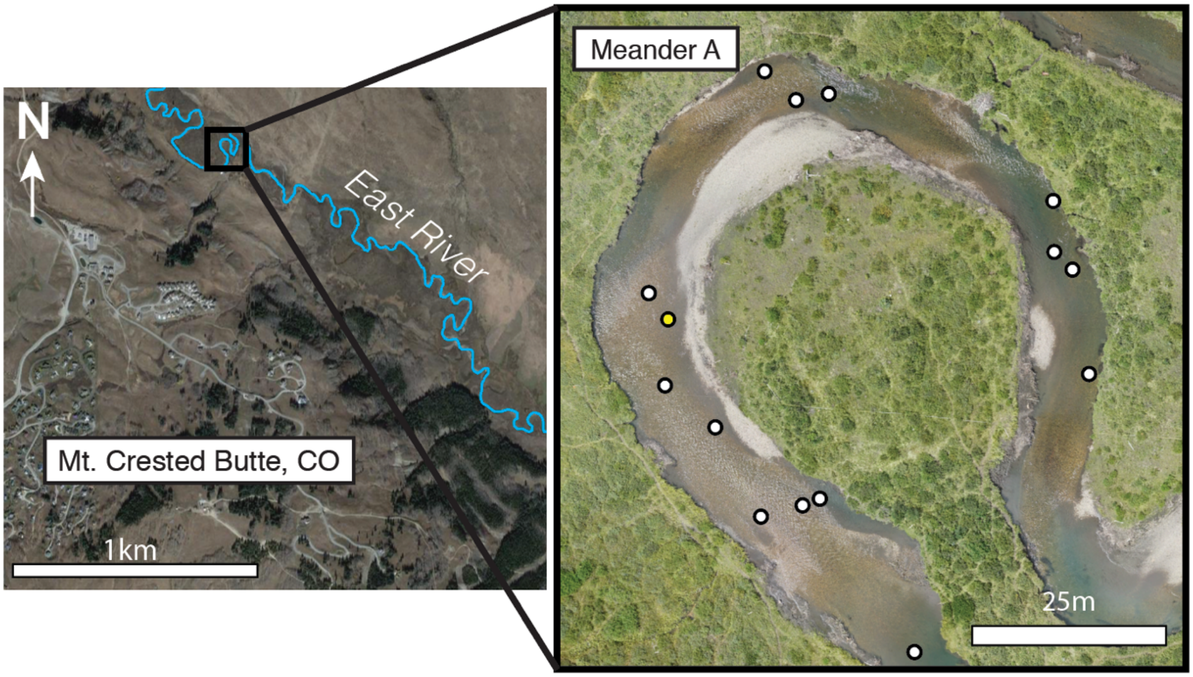


**Figure S1.** Sampling location of surface and pore water samples from Nelson et al. (2019) analyzed here. The sample utilized in Table 1 is highlighted.

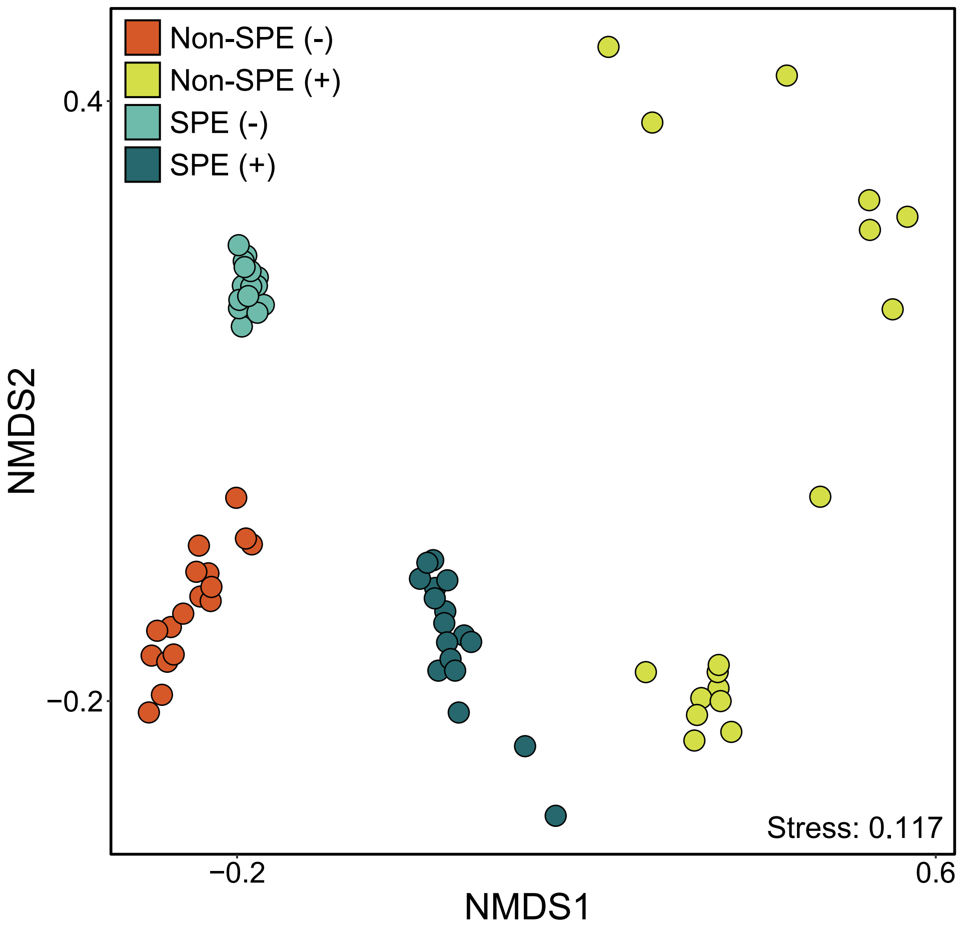

**Figure S2.** Non-metric multidimensional scaling ordination (NMDS) of sample set FTICR-MS data colored by method, indicating clear separation depending on chosen method.

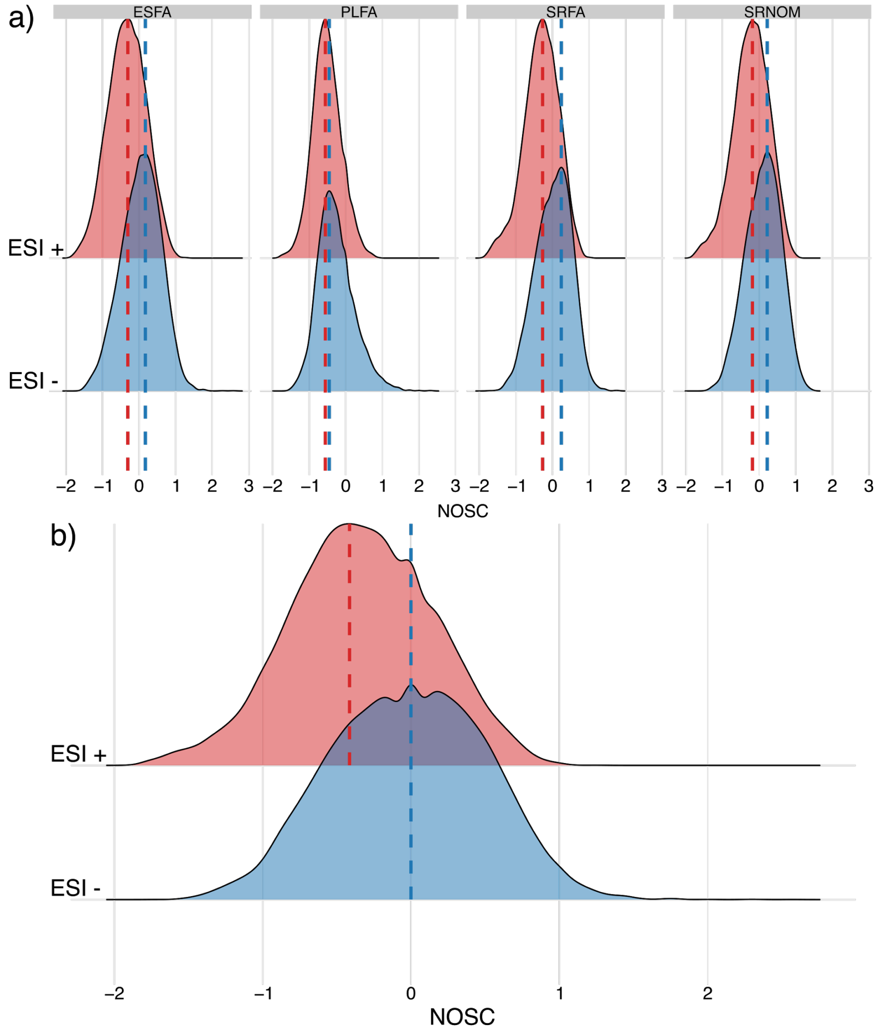


**Figure S3**: NOSC distributions from data obtained from Hawkes et al., 2020. a) NOSC distributions from each sample type (ESFA: Elliot Soil Fulvic Acid, PLFA: Pony Lake Fulvic Acid, SRFA: Suwannee River Fulvic Acid, SRDOM: Suwannee River Natural Organic Matter). b) NOSC distribution for combined sample types. Mann Whitney U and Kolmogorov-Smirnov Tests revealed that ESI – molecular formula had significantly higher NOSC values than ESI + molecular formula.

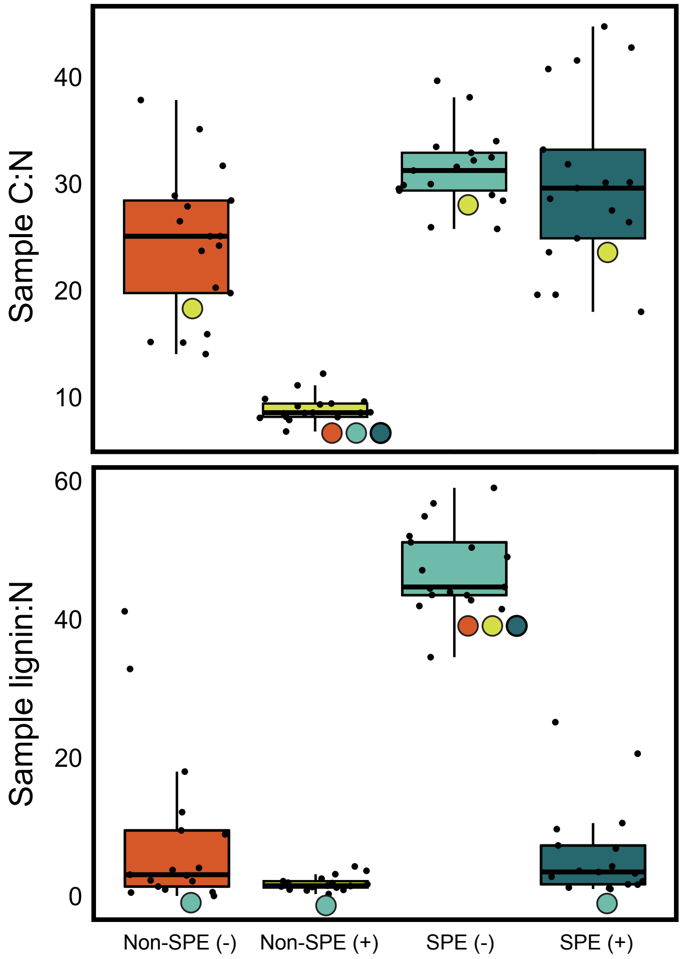


**Figure S4.** The distribution of sample mean C:N and lignin:N across all four FTICR-MS analysis methods. Colored circles indicate significant differences between methods.

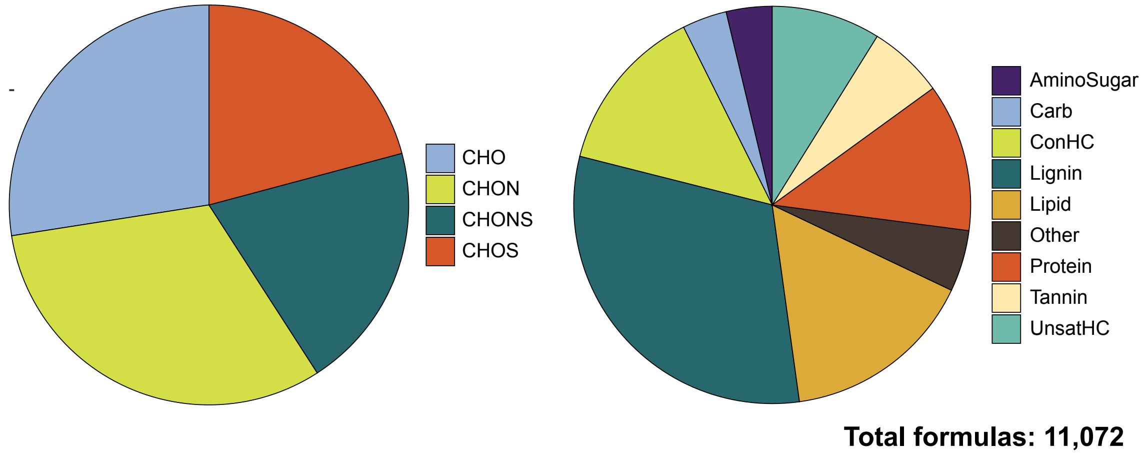


**Figure S5.** Distribution of composition and Van Krevelen classes of 11,072 unique formulas detected with all four methods combined.

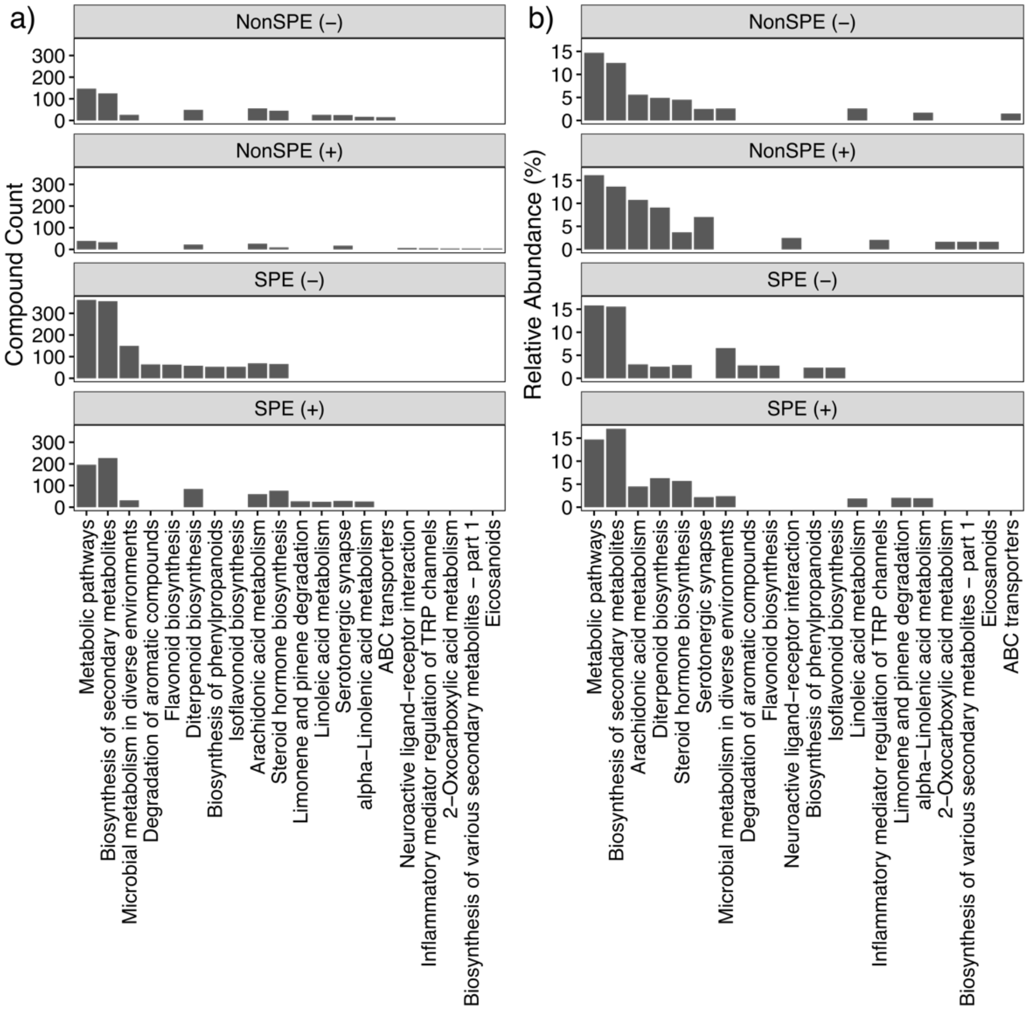


**Figure S7.** Comparison of metabolic pathways observed across treatment methods. a) Absolute compound counts within the 20 most represented pathways (based on actual counts). b) Relative number of compounds observed within the 20 most represented pathways (based on relative abundance).

| Sample name | Sampling depth | Conductivity ($\mu$S) |
|---|---|---|
| A_0_cm | 0 | N/A |
| HRA | 20 | 272.8 |
| 23A | 20 | 289.6 |
| B_40_cm | 40 | 387.4 |
| 49B | 20 | 296.7 |
| 34C | 20 | 376.3 |
| B_10_cm | 10 | 268 |
| 1B | 20 | 283.1 |
| HOBO_10 | 20 | 278.6 |
| HOBO_3 | 20 | 281.4 |
| B_0_cm | 0 | 267.8 |
| HR4 | 20 | 285.6 |
| 36A | 20 | 353.2 |
| HOBO_4 | 20 | 292.7 |
| 47C | 20 | 268.6 |
| 16A | 20 | 269.2 |
| 50C | 20 | 289.8 |

**Table S1.** Sample and corresponding streambed sampling depth and conductivity measurement.

| Treatment | Total Peaks | Filtered Peaks | Assigned Peaks | Unique Formulas |
|---|---|---|---|---|
| Non-SPE (-) | 16384 | 13562 | 1257 | 1184 |
| Non-SPE (+) | 34156 | 30642 | 2773 | 2633 |
| SPE (-) | 20702 | 16508 | 6218 | 5916 |
| SPE (+) | 16586 | 13251 | 3629 | 3299 |

**Table S2.** Overview of the number of peaks from each treatment at various stages of processing. “Total Peaks” are those which were identified in the original dataset, “Filtered Peaks” are those present after using ftmsRanalysis, “Assigned Peaks” are those that were assigned some molecular formula, and “Unique Formulas” are the number of the unique molecular formulas that were assigned.

| Methods | # Common Molecules |
|---|---|
| SPE +/- | 881 (24.2%, 14.1%) |
| Non-SPE +/- | 134 (4.8%, 10.6%) |
| Positive (SPE, Non-SPE) | 511 (14.1%, 18.4%) |
| Negative (SPE, Non-SPE) | 559 (8.9%, 44.4%) |

**Table S3.** Number of molecules shared between indicated methods, with percentages showing the percent of formulas shared from each respective method.
